# Loud noise exposure differentially affects subpopulations of auditory cortex pyramidal cells

**DOI:** 10.1101/2020.08.25.264200

**Authors:** Ingrid Nogueira, Jessica Winne, Thiago Z. Lima, Thawann Malfatti, Richardson N. Leao, Katarina E. Leao

## Abstract

Loud noise-exposure generates tinnitus in both humans and animals. Macroscopic studies show that noise exposure affects the auditory cortex; however, cellular mechanisms of tinnitus generation are unclear. Here we compare membrane properties of layer 5 (L5) pyramidal cells (PCs) of the primary auditory cortex (A1) from control and noise-exposed mice. PCs were previously classified in type A or type B based on connectivity and firing properties. Our analysis based on a logistic regression model predicted that afterhyperpolatization and afterdepolarization following the injection of inward and outward current are enough to predict cell type and these features are preserved after noise trauma. One week after a noise-exposure (4-18kHz, 90dB, 1.5 hr, followed by 1.5hr silence) no passive membrane properties of type A or B PCs were altered but principal component analysis showed greater separation between control/noise-exposure recordings for type A neurons. When comparing individual firing properties, noise exposure differentially affected type A and B PC firing frequency in response to depolarizing current steps. Specifically, type A PCs decreased both initial and steady state firing frequency and type B PCs significantly increased steady state firing frequency following noise exposure. These results show that loud noise can cause distinct effects on type A and B L5 auditory cortex PCs one week following noise exposure. As the type A PC electrophysiological profile is correlated to corticofugal L5 neurons, and type B PCs correlate to contralateral projecting PCs these alterations could partially explain the reorganization of the auditory cortex observed in tinnitus patients.

## Introduction

The auditory cortex is a key structure in generation and maintenance of tinnitus: the perception of a phantom sound in the absence of external stimuli (Elgoyhen et al., 2015). Tinnitus is a prevalent disorder affecting about 15% of the world population (Shargorodsky et al., 2010) and is highly correlated to mood disorders (Langguth et al., 2011; Malouff et al., 2011; Winne et al., 2019). We recently showed that anxiety is prominent in animal models of tinnitus without hearing loss and that tinnitus affect limbic regions related to anxiety/depression (Winne et al., 2020). However, little is known about the after effect of loud noise exposure in auditory cortex neurons. Tinnitus without hearing loss is commonly observed during adolescence and is prevalent in several countries. For example, in the United States, 4,7% of 12-17 years olds reported chronic tinnitus (Mahboubi et al., 2013). Also, a large study of Belgian students found 18.3% of teenagers to report chronic tinnitus but having little concern for being exposed to loud music (Gilles et al., 2013). These numbers are alarming since chronic tinnitus may often correlate with increased stress and depression (Langguth et al., 2011) and can have long term effects on quality of life. As adaptive changes in auditory centers caused by noise may differ considerably between patients with normal hearing thresholds and individuals with hearing loss, this requires further exploration.

Several studies reports adaptive changes in the primary auditory cortex (A1) following acoustic overstimulation of animals (Basura et al., 2015; Munguia et al., 2013; Noreña et al., 2003; Popelár et al., 1987; Seki and Eggermont, 2002; Sun et al., 2008; Takacs et al., 2017). For example, larger evoked potentials and increased firing frequency in the auditory cortex has been observed immediately following noise exposure of rats (Sun et al., 2012). Also, increase in spontaneous firing frequency of the A1 has been shown 4-6 weeks following noise-induced tinnitus in guinea pigs (Basura et al., 2015). Recently noise overexposure was shown to increase the gain of A1 pyramidal neurons projecting to the inferior colliculus and areas of the limbic system (striatum and amygdala) for up to two weeks after the noise exposure (Asokan et al., 2018). Yet, whether loud noise can have different long-term effects on pyramidal neurons with different projection profiles, or whether subtypes of pyramidal neurons are more vulnerable to noise exposure is unknown.

Cortical pyramidal cells (PCs) are heterogeneous in respect to connectivity, as well as morphology and laminar distribution (Harris and Shepherd, 2015; Mason and Larkman, 1990; Wang et al., 2018). Especially heterogeneity among PCs of the main output neocortical layer 5 (L5) have been extensively studied (Hallman et al., 1988; Molnár and Cheung, 2006; Morishima and Kawaguchi, 2006). We have previously used the distinction of type A and type B L5 PCs, where type A PCs correspond large L5 PCs with thick-tufted dendrites to study connectivity between L5 PCs and inhibitory Martinotti cells (Hilscher et al., 2017). Moreover, type A are known to express prominent h-current (Ih) and project subcortically, while type B PCs are thin-tufted, have little Ih and connect contralaterally or to the striatum (Lee et al., 2014). Using patch clamp recordings and principal component analysis, we examine the primary auditory cortex to see if noise exposure of tinnitus-causing noise (Winne et al., 2020) alters membrane or firing properties of type A and type B PCs.

## Methods

### Animals

Wild type mice (c57BL/6) of either sex, age between 2-3 weeks (young) and 5-8 weeks (mature) were used in this study. All experimental procedures followed current guidelines and were approved by the Ethics Committee for the Use of Animals of Federal University of Rio Grande do Norte (CEUA/UFRN) protocol n°097.019/2018. Animals were housed on a 12h/12h day/night cycle and had free access to food and water.

### Noise exposure

Acoustic noise overexposure was carried out in a sound shielded room, inside a sound-isolated cabinet (44×33×24 cm) during the late afternoon. Mice were handled and habituated for 3-5 days by being placed inside an acrylic cylinder (diameter 4×8 cm length), with restraining doors perforated at regular intervals (Acrilart, Natal-RN, Brazil), for 5-10 min at a time. Mice were considered habituated when freely entering the cylinder and there was minimal trace of defecation. A speaker (Selenium Trio ST400) connected to a sound amplifier (Marantz PM8004) and sound board (USBPre2), was placed 10 cm in front of the acrylic cylinder to produce the sound stimulation. The speaker was calibrated using a microphone (Brüel and Kjær 4939-A-011) and adjustment of intensity, frequency and duration of the sound was done using custom written code (Matlab, MathWorks). Awake mice were exposed to broadband noise of 4-18 kHz, at 90 decibel sound pressure level (dBSPL) for 1.5 h, to activate a large portion of the auditory cortex. Immediately following noise overexposure animals were removed from the acrylic cylinder but remained in the sound shielded quiet cabinet, inside a standard plastic cage for another 1.5 hour. This was done since studies have shown that increased ambient noise and acoustic enrichment immediately following a noise trauma can prevent noise-induced tinnitus (Sturm et al., 2017). Following the silence period animals were returned to their home cages in the animal facility for 1 week before being sacrificed for electrophysiological experiments. Control animals were age matched littermates.

### Whole-cell patch clamp

Young mice (P16-23) were sacrificed by decapitation and thereafter immediate brain dissection. To improve cell visibility and cell survival of slices from mature mice (>5 weeks old), mice were routinely perfused prior to slicing and had recovery solution applied (Ting et al., 2014). In detail, mature animals (P38-52) were sacrificed by intraperitoneal injection with ketamine (90 mg/kg) for anesthesia before intracardiac perfusion with artificial cerebrospinal fluid (ACSF) containing (in mM): NaCl, 124; KCl, 3.5; NaH_2_PO_4_, 1.25; MgCl_2_, 1.5; CaCl_2_, 1.5; NaHCO_3_, 30; glucose, 10. Brains were rapidly dissected, the cerebellum and brainstem removed, glued to a platform and submerged in ice-cold sucrose/artificial cerebrospinal fluid (ACSF) consisting of the following (in mM): KCl, 2.49; NaH_2_PO_4_, 1.43; NaHCO_3_, 26; glucose, 10; sucrose, 252; CaCl_2_, 1; MgCl_2_, 4. The brain was cut in coronal slices (300μm thick) using a vibratome (VT1200, Leica, Microsystems) and slices containing the primary auditory cortex were collected and moved to a holding chamber containing normal ASCF, or for >1 month old mice containing N-methyl-D-glucamine (NMDG, recovery) solution (in mM): NMDG, 93; KCl, 2.5; NaH_2_PO_4_, 1.2; NaHCO_3_, 30; HEPES, 20; sodium ascorbate, 5; thiourea, 2; sodium pyruvate, 3; hydrated MgSO_4_; 10; CaCl_2_, 0,5, pH calibrated with HCl to pH 7.3-7.4, following the protocol by (Ting et al., 2014) for 12 minutes to improve cell survival and cell visibility in *in vitro* slices, and next being placed in normal ACSF constantly bubbled with 95% O_2_ and 5% CO_2_ at room temperature (22-24**°**C). Next slices were transferred to a submerged chamber under an upright microscope equipped with differential interference contrast (DIC) optics (Olympus, Japan) and perfused with room temperature oxygenated ASCF (1–1.25 ml/min). Patch pipettes from borosilicate glass capillaries (GC150F-10, Harvard Apparatus, MA, USA) were pulled on a vertical puller (PC-10, Narishige, Japan). Pipette resistances varied from 8-12 MΩ. Pipettes were filled with internal solution containing (in mM): K-gluconate, 130; NaCl, 7; MgCl_2_, 2; ATP, 2; GTP, 0.5; HEPES, 10; EGTA, 0.1 (from Sigma Aldrich, MO, USA). The pH was adjusted to 7.2 using KOH. Whole-cell current clamp recordings were acquired using an Axopatch 200B amplifier (Axon instruments, CA, USA) and digitized with a BNC-2111 panel block (National instruments, TX, USA). Pyramidal cells were visually identified by size and morphology and routinely clamped to −65 mV before breaking in. WinWCP software implemented by Dr.J. Dempster (University of Strathclyde, Glasgow, UK) was used to record electrophysiological signals. Cells with an unstable baseline, membrane resistance and/or more depolarized resting membrane potential than −55 mV was discarded from further analysis.

### Data analysis

Matlab (version 2016a, MathWorks) was used for data analysis of recordings. Resting membrane potential (Vrest) was noted as the baseline in current clamp mode. Membrane resistance was calculated from the current activated by a small test step (5mV, 10ms). Rheobase is the minimum amount of current necessary to generate an action potential (calculated from a ramp protocol from 0 to 200 pA, 500 ms, where the time of the first spike was noted; AP time). Hyperpolarizing sag amplitude was quantified in response to negative current steps (0 to −100 pA, 50 pA decrements, 500 ms) as the difference between peak and steady-state voltage (ΔV mV). The afterdepolarization (ADP) and afterhyperpolarization (AHP) were measured following the termination of a −100pA or +150pA step (500ms duration) respectively, as the peak amplitude subtracted by Vrest. The first AP generated upon positive current injections (ramp from 0 to 200 pA, 500 ms) was analyzed for AP threshold (>10 mV/ms). We also examined the properties of APs using phase-plane plots, which show the derivative of membrane potential (dVm/dt) as a function of instantaneous membrane potential. Phase plots were obtained by plotting dV_m (obtained using the matlab command *diff*) vs V_m.top). Firing frequency was analysed from AP generated by depolarizing current injections (50 to 400 pA, 50 pA increments, 1s duration). Initial frequency denotes the frequency of the first two APs, calculated as the inverse of the first interspike interval (ISI). Steady-state frequency denotes the frequency of the last 3 APs, calculated as the inverse of the mean of the last three interspike intervals. The initial and steady-state gain was calculated by fitting a trendline to both initial and steady state frequency in response to 150, 200 and 250 pA current steps and quantifying the slope (Hz/pA).

### Statistical Analysis

Statistical analysis of the predictability of cell type and effect of condition, the experiment followed a 2^2^ factorial design, hence with 2 factors: cell type and noise-exposure (experimental condition), both with 2 levels. Sample sizes (after outliers removal) were 11(8), 13(8), 19(17), 21(19) for groups A-control, B-control, A-noise, B-noise, respectively. Principal component analysis (PCA) was computed from the correlation matrix of quantitative variables of the dataset, namely: absolute sag, ADP and AHP, resting potential, input resistance, AP threshold, AP time, rheobase, initial ISI, initial frequency, steady-state ISI and frequency, initial and steady state frequency-current gain. The amount of information accounted by each principal component (PrC) is measured by its relative variance as compared to total variance. The meaning of each PrC was interpreted according to the signal and absolute value of the coefficients of the linear combination that generates it. The model for cell type classification was developed by means of logistic regression. When separated according to cell type only, all variables were shown to be compatible with the normal distribution. The selection of variables to the model was assisted by forward and backward stepwise selection and highly correlated variables were avoided. The model was defined based on the lowest deviance and AIC (Akaike information criteria) obtained. To perform parametric inference, 8 outliers were discarded because they were hampering the normalization of at least one variable. The variables incompatible with the normal distribution were transformed by injective functions: Box-Cox transform, Ln or inverse. Then, the effect of factors was tested by multivariate analysis of variance (MANOVA). To further identify response variables that most contribute for each effect, individual 2-way ANOVA was used as post-hoc. For those untransformed variables, we also estimated effect sizes and an empirical model, which was based on multiple linear regression. For both ANOVA and linear regression, normality, independence, and variance homogeneity assumptions were checked by the analysis of the residues. Main effects were computed as the mean difference between levels of correspondent factor, whereas interaction effect was estimated by half the difference of the effect of one factor relative to both levels of the other factor. For all hypothesis tests, a significance level of 0.05 was used. For basic comparison between type A and type B variables in tables, two-way Student’s *t*-test, equal variance was applied, and data reported as standard error of the mean (s.e.m).

### Injection of genetically encoded calcium indicator

For L5 auditory cortex calcium imaging, mice were injected with the virus encoding AAV9.CaMKII.GCaMP6f (pENN.AAV.CamKII.GCaMP6f.WPRE.SV40, gift from James M. Wilson, Addgene viral prep # 100834-AAV9) at the following coordinates: AP −2.18 mm, ML +3.35 mm, DV 2.10 mm; and AP −2.90 mm, ML +4.20 mm, DV 2.25 mm – 0,75μl per site. Ten days after the injection, the mice underwent another surgery for implanting the prism/gradient-index (GRIN) lens, as previously (Gulati et al., 2017). The prism/GRIN (length total: 3.68 mm; length prism: 1mm; diameter: 1.0 mm; pitch: 0.23; object working distance: 0.3 mm, Go Foton Corporation) was implanted in the coordinates: – 2.70 mm AP, +3.95 mm ML, 2.70 mm DV. Two weeks later, mice were checked for GCaMP expression in the tissue through the prism probe. Next, a second GRIN lens, a relay GRIN lens (length: 4.31mm; diameter: 1.8 mm Edmond Optics) was attached in the prism probe and a base plate attached in the skull of each animal, in the optimum image plane. The prism measured 2×2mm and was glued to a 2mm diameter GRIN lens. A 2mm grin lens/prism was also tested with non-optimal results. The GRIN lens/prism projected the image in a second grin lens placed 0.4mm above (air).

### Calcium imaging and analysis

At the beginning of each imaging session, the animals were anesthetized with isoflurane 5% to fix the miniature microendoscope (Miniscope V3, UCLA) (Aharoni and Hoogland, 2019) in the base plate, and 30 minutes after the animals recovery from anesthesia the imaging session started. Focus of the miniscope was adjusted to an imaging field about 300 μm away from the prism. Optimum laser power (20% of the maximum), imaging gain, and focal distance were selected for each animal and conserved across all sections. The Ca2+ signal was acquired at a frame rate of 30 frames per seconds, using the Miniscope controller software (http://www.miniscope.org). Two consecutive days prior to the first recording session, the mice were habituated to the miniscope and the experimental room for 20 minutes. To generate cortical activity in the auditory cortex animals were exposed to stimuli consisting of narrow band sounds at frequencies of 2–20 kHz ±0.5KHz, 60db amplitude. Noise trauma was performed as described above and the cortex was imaged again 1 week after the trauma. Calcium activity was extracted using constrained non-negative matrix factorization (Pnevmatikakis et al., 2016) implemented in the Minian package from Denise Cai’s lab (https://github.com/DeniseCaiLab/minian). In order to find the same cells across different experimental sessions (Sheintuch et al., 2017), neurons were tracked using the Cell Registration Matlab package (https://github.com/zivlab/CellReg).

## Results

To test whether noise overexposure (4-18kHz at 90 dBSPL for 1.5 hr, 1.5 hr silence post noise exposure) affects L5 PCs, we performed current clamp recordings of pyramidal cells (n=87 cells) in layer 5 of the primary auditory cortex 1 week following noise exposure (Figure 1A). We first established that mice >5 weeks old showed mature membrane properties for both subtypes of PCs examined compared to slices from 2-3 week old mice (Supplementary figure 1, Supplementary table 1). PC subtype was identified post hoc by fitting a model for cell type classification based on logistic regression (Table 1).

**Table 1.** Logistic regression model for pyramidal cell type classification.

|  |  |  |
| --- | --- | --- |
| Model: | $P("A\ cell") = \frac{e^{-300.16 + 89.98\ Abs.ADP + 43.68\ Abs.AHP}}{1 + e^{-300.16 + 89.98\ Abs.ADP + 43.68\ Abs.AHP}}$ | |
| AIC: | 6 | Predictive power = 1 (accuracy: 64/64) |
|  | Deviance | Degrees of freedom |
| Null | 88.473 | 63 |
| Residual | $2.7126 \times 10^{-10}$ | 61 |

Akaike Information Criteria (AIC), Absolute afterdepolarization (Abs.ADP), Absolute afterhyperpolarization (Abs.AHP), P("A cell") - probability of a cell to be classified as type A

**Figure 1.**
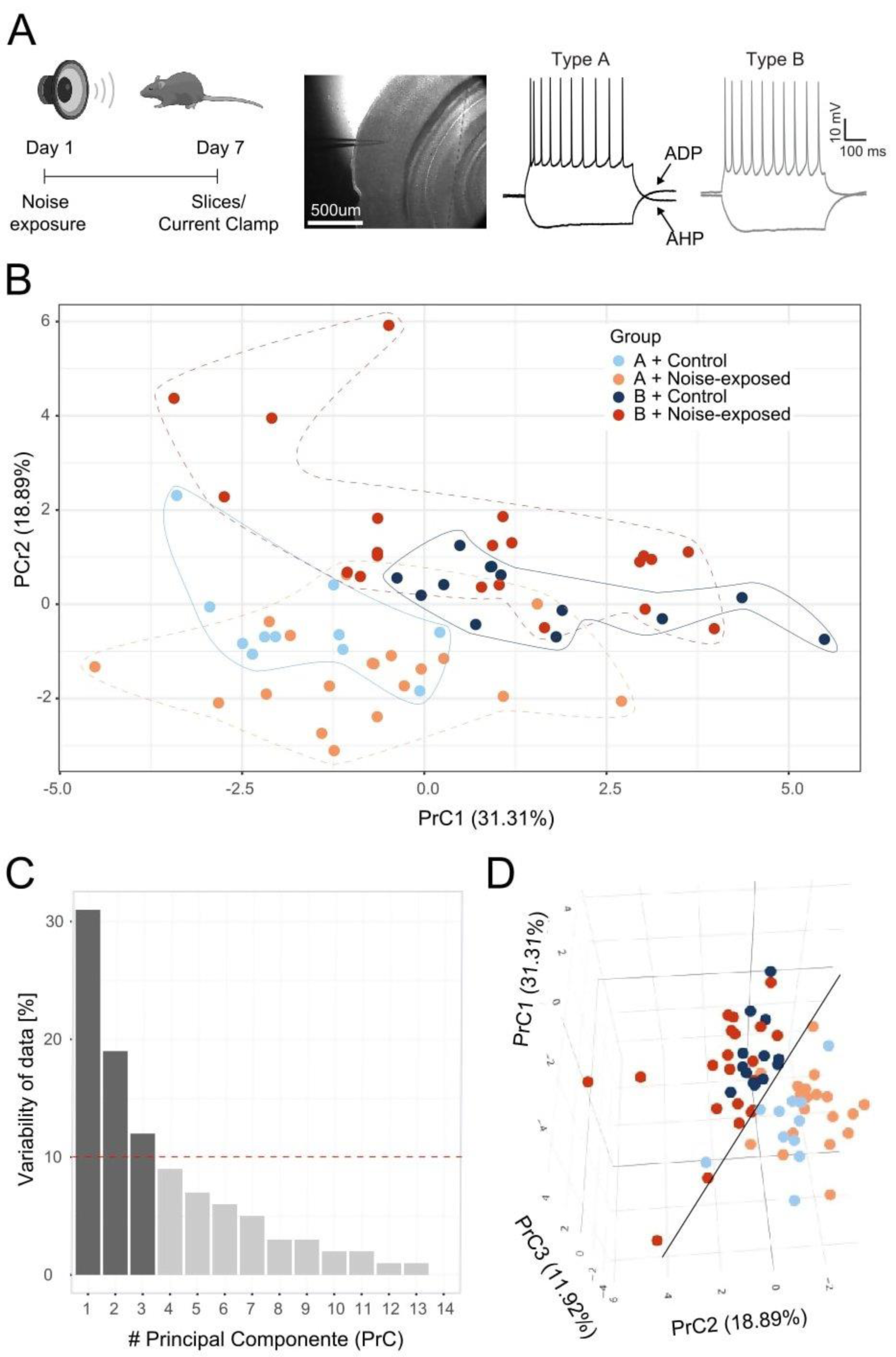
Principal component (PrC) analysis of PC parameters. (A) Schematic representation of the experiment. *Left*: timeline of the acoustic noise trauma and electrophysiological recordings. *Middle*: brightfield image of the primary auditory cortex with the pipette pointing towards layer 5. *Right*: Representative traces of control type A and B PCs in response to −100 and a 150 pA steps. (B) Visual clusterization of the data based on the first two PrC accounting for 50.2% of whole dataset information. Type A control (light blue), type B control (dark blue), type A noise-exposed (light beige) and type B noise-exposed (red). Clusters of distribution of cells are highlighted with lines contouring the corresponding data points. (C) The relevance of each PrC, by the proportion of the whole dataset information relating to each PrC. The red dotted line highlights 10% cut-off for useful PrCs. (D) The inclusion of the third PrC adds 11.92% of whole dataset information to the visual clusterization. The black line highlights the separation of type B (upper in the plot) and type A (lower in the plot) pyramidal cells.

These results showed that afterdepolarization (ADP) and afterhyperpolarization (AHP) were sufficient for cell classification (maximum predictive power of the model) (Gee et al., 2012; Hilscher et al., 2017; Joshi et al., 2015; Lee et al., 2014). Thereby, L5 PCs are hereafter referred to as type A or type B PCs (Lee et al., 2014). Next, principal component analysis based on 11 electrophysiological parameters (Table 2) from control and noise overexposed mice was carried out to illustrate the variability of the data from type A and type B PCs from control and noise exposed mice (Figure 1). The values of the first two principal components (PrC1-2, Supplementary table 2) corresponded to 50% of the dataset (Table 2) information (i.e., variability, PrC1: 31%, PrC2: 19%, Figure 1B-C).

**Table 2.**
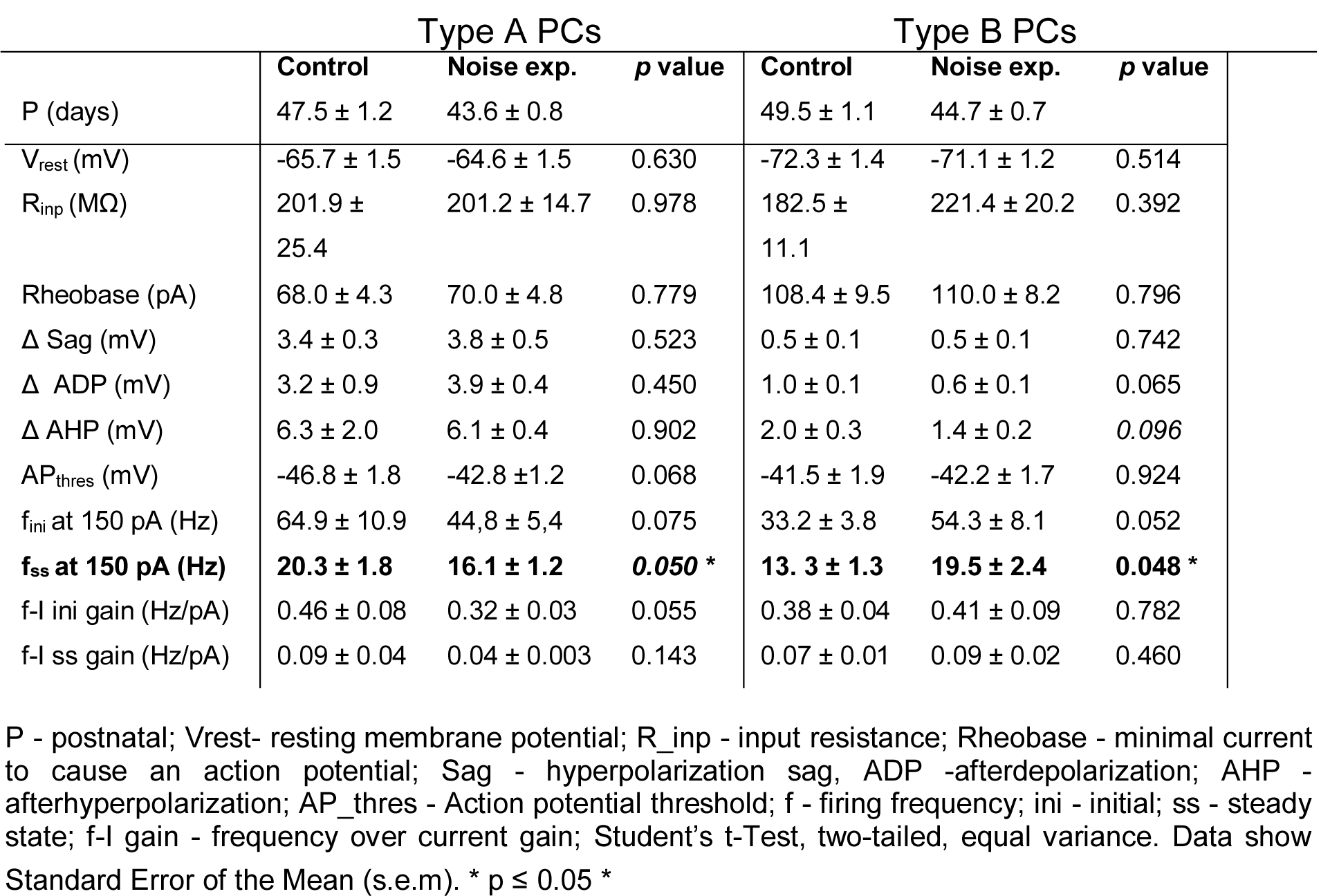
Type A and type B pyramidal cells show different firing frequency following noise-overexposure.

|  | Type A PCs |  |  | Type B PCs |  |  |
| --- | --- | --- | --- | --- | --- | --- |
|  | Control | Noise exp. | p value | Control | Noise exp. | p value |
| P (days) | 47.5 ± 1.2 | 43.6 ± 0.8 |  | 49.5 ± 1.1 | 44.7 ± 0.7 |  |
| V<sub>rest</sub> (mV) | -65.7 ± 1.5 | -64.6 ± 1.5 | 0.630 | -72.3 ± 1.4 | -71.1 ± 1.2 | 0.514 |
| R<sub>inp</sub> (MΩ) | 201.9 ± 25.4 | 201.2 ± 14.7 | 0.978 | 182.5 ± 11.1 | 221.4 ± 20.2 | 0.392 |
| Rheobase (pA) | 68.0 ± 4.3 | 70.0 ± 4.8 | 0.779 | 108.4 ± 9.5 | 110.0 ± 8.2 | 0.796 |
| Δ Sag (mV) | 3.4 ± 0.3 | 3.8 ± 0.5 | 0.523 | 0.5 ± 0.1 | 0.5 ± 0.1 | 0.742 |
| Δ ADP (mV) | 3.2 ± 0.9 | 3.9 ± 0.4 | 0.450 | 1.0 ± 0.1 | 0.6 ± 0.1 | 0.065 |
| Δ AHP (mV) | 6.3 ± 2.0 | 6.1 ± 0.4 | 0.902 | 2.0 ± 0.3 | 1.4 ± 0.2 | 0.096 |
| AP<sub>thres</sub> (mV) | -46.8 ± 1.8 | -42.8 ± 1.2 | 0.068 | -41.5 ± 1.9 | -42.2 ± 1.7 | 0.924 |
| f<sub>ini</sub> at 150 pA (Hz) | 64.9 ± 10.9 | 44.8 ± 5.4 | 0.075 | 33.2 ± 3.8 | 54.3 ± 8.1 | 0.052 |
| **f<sub>ss</sub> at 150 pA (Hz)** | **20.3 ± 1.8** | **16.1 ± 1.2** | **0.050 \*** | **13.3 ± 1.3** | **19.5 ± 2.4** | **0.048 \*** |
| f-I ini gain (Hz/pA) | 0.46 ± 0.08 | 0.32 ± 0.03 | 0.055 | 0.38 ± 0.04 | 0.41 ± 0.09 | 0.782 |
| f-I ss gain (Hz/pA) | 0.09 ± 0.04 | 0.04 ± 0.003 | 0.143 | 0.07 ± 0.01 | 0.09 ± 0.02 | 0.460 |

P - postnatal; V<sub>rest</sub>- resting membrane potential; R<sub>inp</sub> - input resistance; Rheobase - minimal current to cause an action potential; Sag - hyperpolarization sag, ADP -afterdepolarization; AHP - afterhyperpolarization; AP<sub>thres</sub> - Action potential threshold; f - firing frequency; ini - initial; ss - steady state; f-I gain - frequency over current gain; Student's t-Test, two-tailed, equal variance. Data show Standard Error of the Mean (s.e.m). \* p ≤ 0.05 \*

Plotting the first and second principal components showed a minimal overlap between the type A and type B PC clusters, where type B cells (exhibited in dark colors) gather at greater values of the PrC2 than type A cells (Figure 1B). Pyramidal neurons from control condition cluster at intermediary values of PrC2 while noise-exposed subjects spread in the opposite direction depending on type A (downwards) and type B (upwards) values. The first PrC was mostly relevant for separating cell types in control condition (Figure 1B). Addition of the third PrC (accounting for an additional 12% of variability, Figure 1C) allowed for a more comprehensive representation of the data in a 3D plot preserving 62% of the whole dataset information (Figure 1D). In the 3D plot the distinction between subtype of PCs is visible (black line) and for type A PCs the experimental condition became more evident (separation of light blue and light pink clusters, Figure 1D). For type B cells, control (dark blue) and experimental conditions (dark red) remain overlapped, but noise exposed type B PCs spread more than the control type B PCs cluster (Figure 1D). A multivariate analysis of variance (MANOVA) test confirmed that cell type had a significant effect (p = 1.58 × 10^−11^) on response variables and such effect was different in control and noise-exposed animals (Supplementary table 3). This shows that type A and type B PCs are different electrophysiologically and that noise exposure increases variability but in different directions for type A and type B PCs.

Examining each electrophysiological parameter separately we first show that noise exposure did not alter hyperpolarization sag, ADP and AHP in responses to a negative (−100 pA, 500 ms) and a positive (150 pA, 500 ms) current step for L5 type A or type B PCs from noise-exposed or control mice (Figure 2). In both conditions, type A PCs (n=30) had pronounced sag, ADP and AHP compared to type B PCs (n=34) that showed generally flat response following negative and positive current injections (Figure 2A). Differences between type A and type B PCs were equally recognizable following noise exposure (Figure 2B) with a >6 mV cut-off criteria of the sum of sag, ADP and AHP amplitude (Figure 2C), similar to previously shown (Lee et al., 2014).

**Figure 2.**
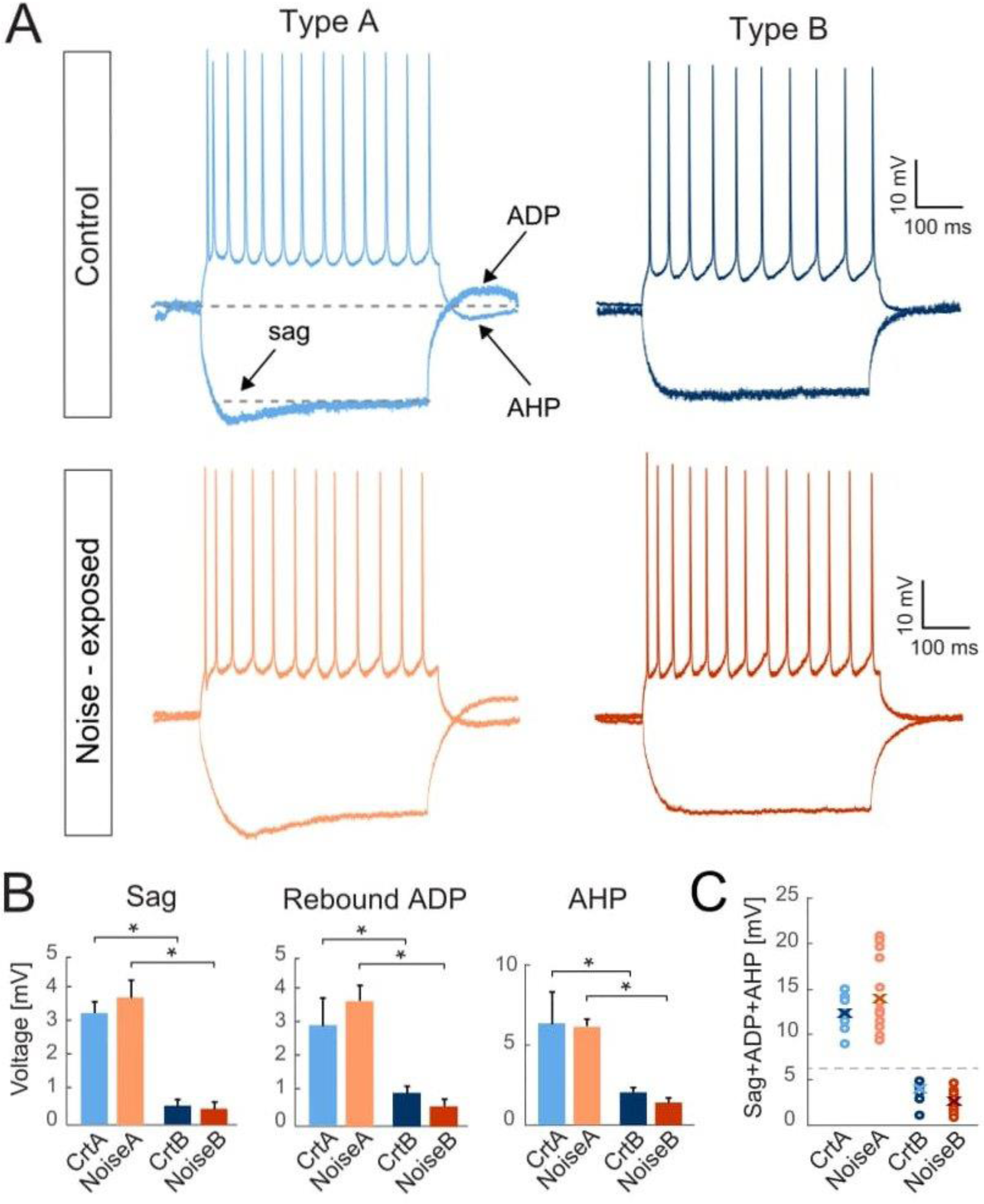
Criteria for separating L5 PCs into type A and type B remains robust between control and noise-exposed experimental groups. (A) Representative traces in response to −100 pA and 150 pA steps from L5 type A and type B PCs from control (*up*, blue) and noise-exposed (*bottom*, red) mice. (B) Type A and type B cells show distinct values for, hyperpolarizing sag, rebound afterdepolarization (ADP) and afterhyperpolarization (AHP) potential for control and noise-exposed groups. Error bars – s.e.m., Student’s *t-*test, two tailed, equal variances.

Passive membrane properties (see Table 2) of L5 PCs from control and noise exposed mice showed that type B PCs have a more hyperpolarized membrane potential than type A PCs but that resting membrane potential and input resistance are not altered for type A and type B PCs following noise exposure. Examining action potential properties showed no difference in action potential threshold, although type A PCs showed a trend towards a slightly depolarized AP threshold in cells from noise-exposed animals (ctr A: −46.8 ± 1.8 mV vs. noise A: −42.8 ± 1.2 mV, p= 0.068) while AP threshold was robust for type B PCs from the two groups (ctr B: −41.5 ± 1.9 mV vs. noise B: −42.2 ± 1.7 mV, p=0.924). Therefore, we also carried out phase plot analysis (Trombin et al., 2011) of the action potential for control and noise-overexposed type A and type B L5 PCs (Figure 3A). As described in Trombin et al., 2011, the first derivative of the AP is represented as a loop highlighting the threshold membrane potential (*V*_thres_), and the maximal voltage amplitude (*V*_max_), with the depolarization and repolarization phases (slopes) characterized as the upper and lower portions of the loop, respectively (Trombin et al., 2011) (Figure 3B *left*). The phase plots did not reveal any differences of type A or type B PCs following noise exposure (Figure 3B). The slope of repolarization (Srepol; ctr A: 3.9 ± 0.3 mV/ms vs. noise A: 4.7 ± 0.25 mV/ms, p=0.09, and, ctr B: 4.3 ± 0.2 mV/ms vs. noise B: 4.4 ± 0.3 mV/ms. p=0.63) was not different following noise exposure. Neither was slope depolarization (Sdepol; ctr A: 1.35 ± 0.2 mV/ms vs. noise A: 1.4 ± 0.08 mV/ms, p=0.66, and, ctr B: 1.4 ± 0.08 mV/ms vs. noise B: 1.5 ± 0.09 mV/ms, p=0.29) or Vmax and repolarization voltage (Figure 3B) showing that noise overexposure does not alter the shape of the first action potential of L5 type A and type B PCs.

**Figure 3.**
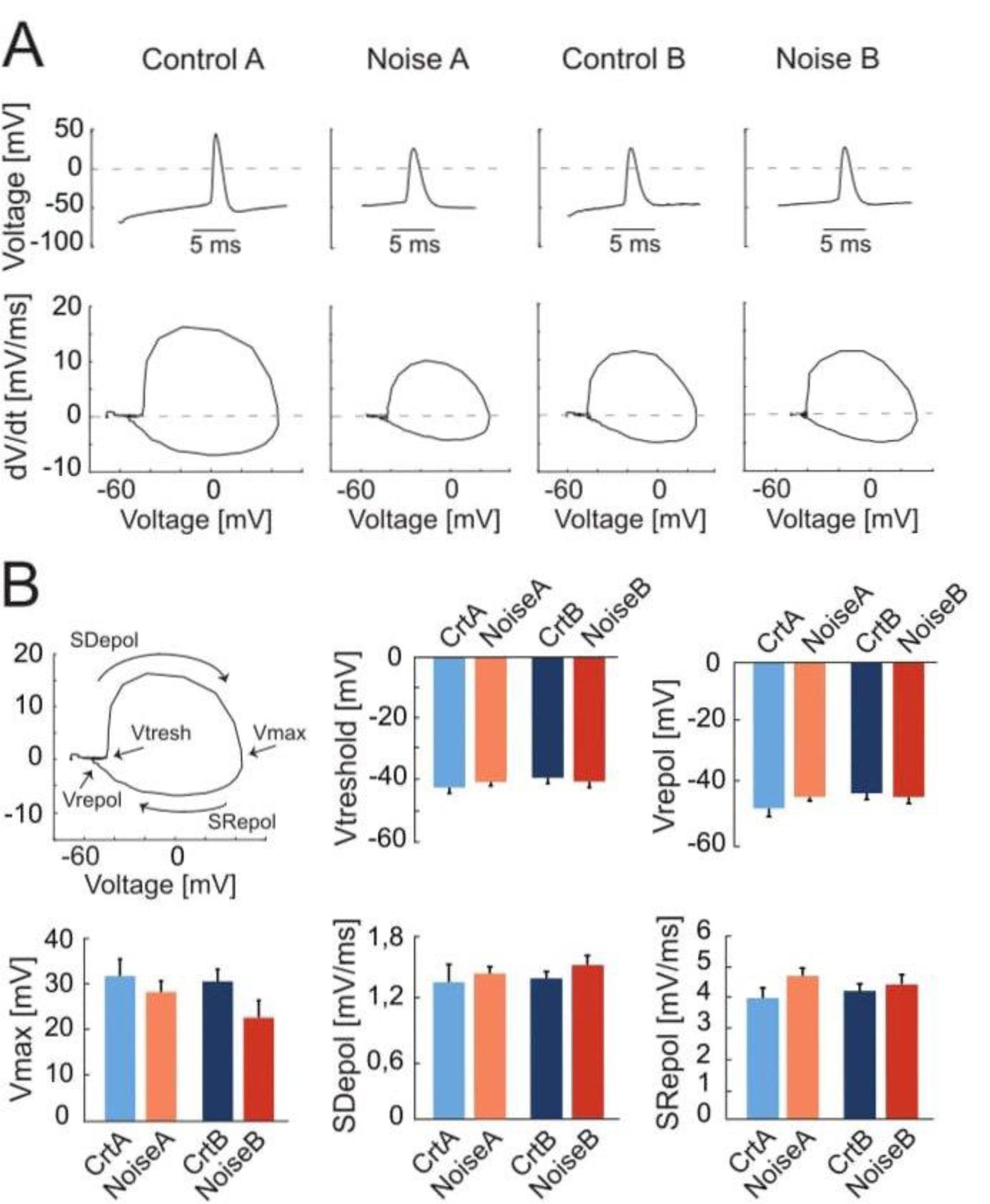
Phase plot analysis of Action Potentials from L5 PCs. (A) Representative traces of first APs recorded from control and noise-overexposed type A and B PCs in response to a 150 pA current injection (*up*) and the relative phase plots (*bottom*) showing depolarization and repolarization phases. (B) The first panel shows a phase plot and the representation of the threshold membrane potential (Vtresh), the maximal voltage peak of the AP (Vmax), the repolarization potential (Vrepol), and the upper and lower portions of the loop which represents the depolarization and repolarization phases (Slopes), respectively. Bar graphs of Vtreshold, Vmax, Vrepol, Sdepol and Srepol for type A and type B PCs from control and noise-exposed mice. Error bars – s.e.m., Student’s *t-*test, two tailed, equal variances.

Next, firing frequency in response to depolarizing current steps (100-250 pA, 1s duration) was examined for type A and type B L5 PCs from control and noise exposed mice (Figure 4). We found that the average initial firing frequency (from the ISI of the first two APs) was significantly lower for type A PCs one week after noise exposure (Figure 4C-D). Type B PCs showed no difference in initial firing frequency in slices from control or noise exposed mice (Figure 4, see Table 2). The initial frequency increased linearly in response to 1s duration current steps (100-250 pA) for both type A and type B PCs from both experimental conditions (Figure 4C, *left*). Noise exposure thereby appears to decrease the initial firing frequency (or initial doublet frequency) of type A PCs and make initial frequency more similar to type B PC initial frequency (Figure 4C). Next, comparing steady-state firing frequency (last three APs in response to +150pA step, 1 s duration) also showed a decrease in frequency for type A PCs from noise exposed mice (ctr A: 20.3 ± 1.8 Hz, n = 11 vs. noise A: 16.1 ± 1.2, n = 19, Hz, *p* = 0.050). On the contrary, type B PCs showed an increased average steady state firing frequency from noise exposed mice (ctr B: 13.3 ± 1.3 Hz, n = 13 vs. noise B: 19.5 ± 2.4, n = 21, *p* = 0.048, Figure 4A-D). Specifically, type B PCs from noise-exposed animals showed a more pronounced increase in firing frequency in response to increasing current injections (150-250pA) compared to type A PCs and control type B PCs (Figure 4C, *right*).

**Figure 4.**
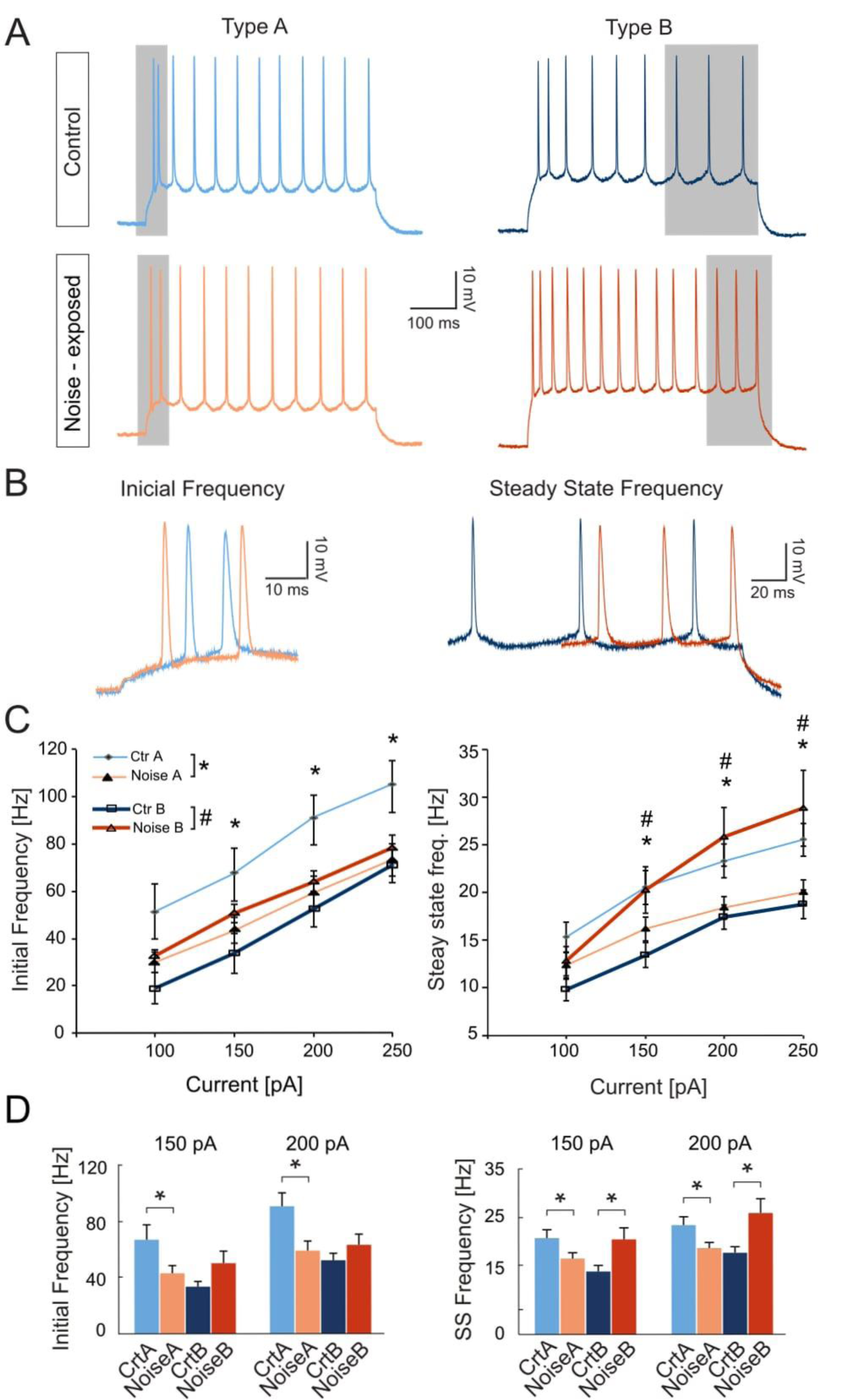
Noise exposure alters steady state frequency in opposite directions for L5 type A and type B PCs. (A) Representative traces illustrating firing frequency in response to a 1s current injection of 150 pA. Gray shadows highlight the first two APs used for calculating initial frequency and the last 3 APs used for calculating steady-state frequency. (B) Higher magnification of traces highlighting the difference in initial firing frequency of L5 type A PCs from control and noise exposed mice (*left*), and the difference in steady state firing for control and noise-exposed L5 type B PCs. (C) Initial frequency over current plots shows a decrease in initial frequency for type A PCs after the noise overexposure (*left*). *Right:* steady state frequency-current plot shows significant increase in steady state frequency for type B PCs after noise exposure while type A cells on the contrary shows a decrease in steady state frequency. (D) Bar graphs showing initial and steady state firing frequency in response to a 150 pA and 200pA stimulation. Error bars – s.e.m., Student’s *t-*test, two tailed, equal variances. (**\***) denotes p<0.05 for type A control vs type A noise-exposure, (**#)** denotes p≤0.05 for type B control vs. type B noise-exposure. (see Table 2 for specific values).

To further investigate differences in firing frequency we examined frequency-current (f-I) gain by comparing the average slope (Hz/pA) of initial and steady-state frequency in response to depolarizing current steps (150, 200, 250pA) of type A and type B L5 PCs (Figure 5A-B). Comparing the average slope in initial frequency shows a trend of type A PCs decreasing gain following noise exposure (ctr A: 0.46 ± 0.08 Hz/pA vs. noise A: 0.32 ± 0.03 Hz/pA, *p* = 0.055) while type B PCs showed no difference in initial firing frequency gain. For the average steady state frequency gain there was no difference between control and noise-exposed cells for type A or type B PCs (Table 2, Figure 5C). However, the differences in steady state gain becomes significantly different between type A and type B PCs following noise exposure (noise A: 0.04 ± 0.0 vs. noise B: 0.09 ± 0.02, *p* = 0.03, Figure 5C, right). This supports that noise-exposure has long term effects on firing frequency of type A and B PCs in the opposite direction.

**Figure 5.**
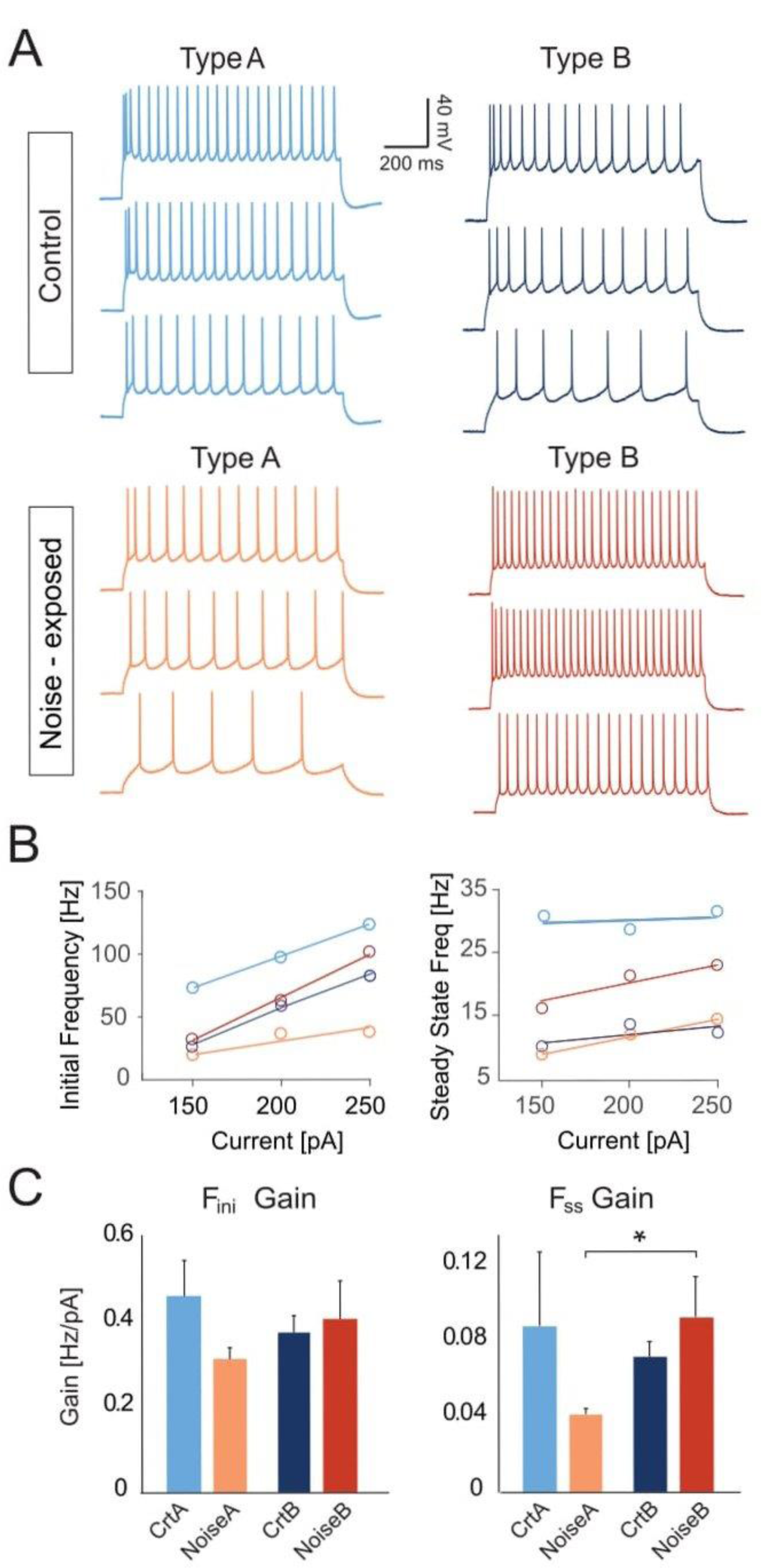
Noise exposure increases frequency over current gain of type B PCs. (A) Representative current clamp traces in response to 150pA, 200pA and 250pA current injections (1 s duration) for type A and type B PCs from control (*up*) and noise-exposed (*down*) mice. (B) Example of a f-I plot showing initial frequency vs current injection (*left*) and steady state frequency vs current injection (*right*) for the neurons shown in ‘A’. (C) Bar graphs of gain (frequency/current) of initial firing frequency (*left*) and steady-state frequency (*right*) for L5 type A and type B PCs from control and noise-exposed mice. Error bars – s.e.m., Student’s *t-*test, two tailed, equal variances.

Finally, in order to test if emerging single photon calcium imaging technology could be used to assess changes in L5 neurons caused by noise trauma (or other tinnitus induction methods) we performed a pilot imaging experiment in one mouse before and after noise exposure. These experiments were not intended to produce statistically significant *in vivo* data in L5 PCs but rather to show the potential use of the technology and design a framework for calcium imaging experiments in tinnitus. A custom 2mm GRIN lens coupled to a prism was constructed to image L5 PCs perpendicularly (Figure 6A). CaMKII-GCamp6f expression was mostly observed in L5 PCs (Figure 6A). In the initial session (before noise exposure), 60 individual neurons were detected and 60 after noise exposure (9 days later: 2 days before noise-exposure and 7 days after) (Figure 6B). Out of the 60, 38 cells were identified in both sessions (Figure 6B). Response to sound stimulation among these cells was not altered between the two sessions but 12 cells increased, and 14 cells decreased non-stimulus related firing (Figure 6C).

**Figure 6.**
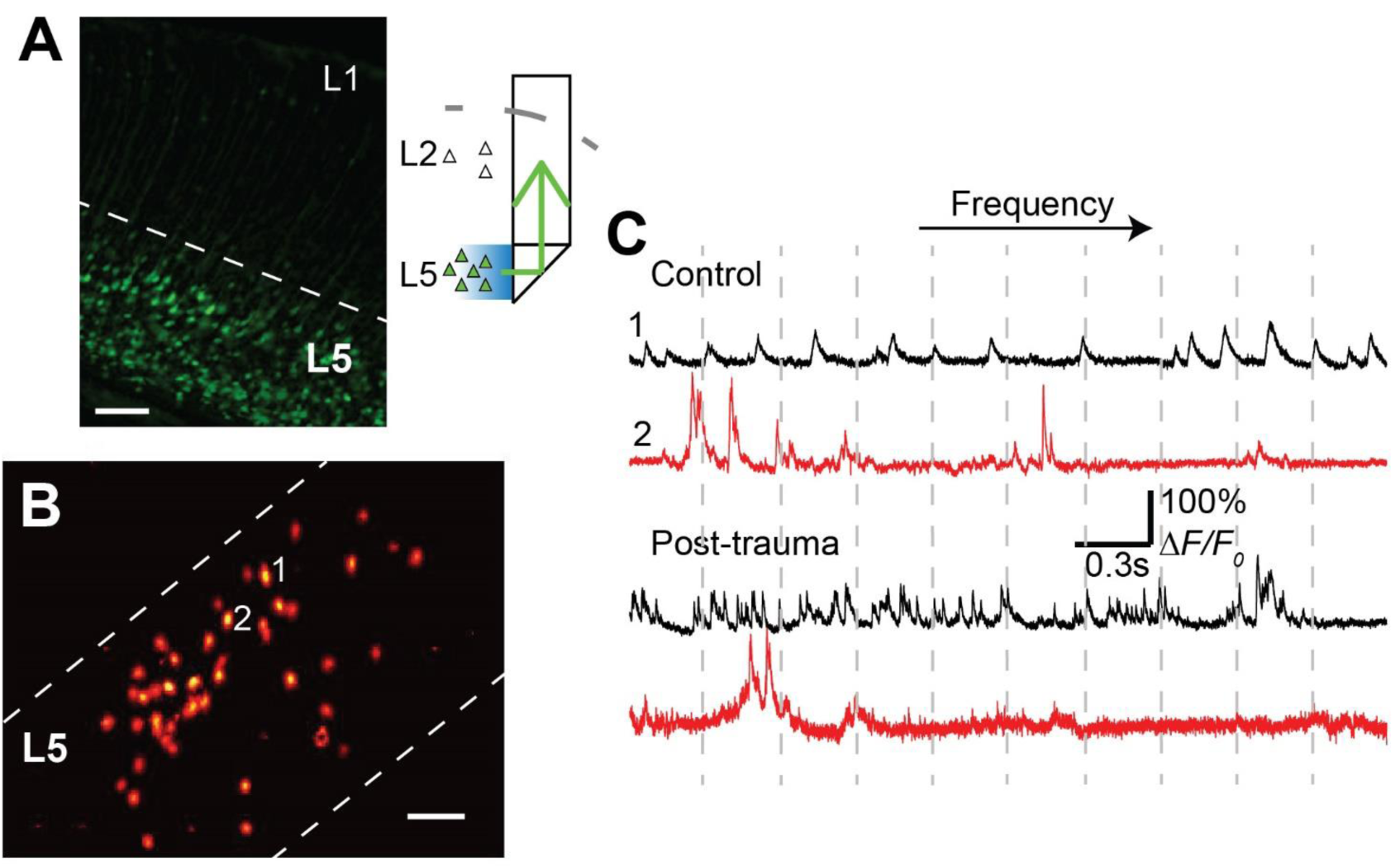
Example of L5 PC cell response to narrow band sound tones before and after noise exposure. (A) Photomicrography showing GCamp6f expression (CamKIIa promoter) in the auditory cortex. *Right*. Diagram showing how imaging of L5 neurons was achieved. (B) Neurons highlighted using the non-negative matrix factorization algorithm. (C) Example of calcium activity of two neurons before and after noise exposure. The traces shown are in response to 60db sound stimulation to narrow band (±0.5KHz) sounds of frequencies ranging from 2 to 20KHz (2KHz steps in every 30s).

## Discussion

In this work, we show that noise-exposure has dissimilar effects on firing frequency of two main types of L5 PCs of the primary auditory cortex of mice. Layer 5 pyramidal cells can be divided into many subtypes based on their projections but for simplification L5 PCs are often grouped into two main subtypes: commissural and corticofugal-projecting PCs (Oswald et al., 2013). Here we investigate those main subtypes but based on electrophysiological profiles. We found that after noise exposure, type A and B cells are still classifiable by PCA; however, only clusters for type A cells are further distinguishable between control and noise exposure groups. We also found that, after noise exposure, type A PCs lowered initial and steady state firing frequency while type B PCs only showed increased steady state frequency. Our results show that loud noise can cause distinct effects on type A and B L5 auditory cortex PCs one week following noise exposure.

Our whole-cell patch clamp experiments found that firing of L5 type A and B PC was differentially affected by a single previous session of noise exposure. Interestingly, our *in vivo* pilot L5 PC calcium imaging experiment also found cells with increased and decreased firing frequency after noise exposure (Fig. 6C). Miniscope imaging allows for identifying individual neurons over time and thus makes it possible to compare firing of individual PCs both before and one week after noise exposure, something that could revolutionize tinnitus research. However, additional tracers are needed to specify subtypes of L5 PCs identified by miniscope imaging. In general, type A and B PCs are broad classes of PCs that differ in morphology and response to hyperpolarization. Subcortical/corticofugal projecting L5 PCs show a thick-tufted dendritic tree, prominent hyperpolarization-activated current (Ih) (i.e., type A PCs), while contralateral/callusal or striatially projecting PCs show thin-tufted dendrites and have less Ih (i.e., type B) (Lee et al., 2014; Oswald et al., 2013). Here, we did not find any effect of noise exposure on h-current (as indicated by sag size). Instead, voltage clamp data suggest that noise exposure may affect outward currents that control firing frequency (Leão et al., 2010). For example, one hour of sound stimulation quickly increases high-threshold voltage dependent potassium channel expression (responsible for sustaining high frequency firing) in brain stem auditory neurons (Leão et al., 2010). Future studies should aim to investigate how specific outward currents are affected by noise trauma, especially as potassium channel modulators are used in tinnitus treatment (Sun et al., 2015).

In our study we used principal component analysis for illustrating changes of electrophysiological parameters following noise exposure. Principal components are linear combinations of original variables and their meaning rises from directions (negative or positive) and magnitude of corresponding coefficients that generate each PrC (see Supplementary Table 2). In this work, the first PrC can be regarded as an inertia of the PC to fire action potentials, due to higher values of rheobase, initial ISI, AP threshold and timing of first AP, all leading to higher values of PrC1. In the same way, lower values of initial and steady state frequency and resting potential also generate higher PrC1 values. Next, PrC2 may be interpreted as cells with higher values of steady-state frequency, f-I gain for steady state frequency, rheobase, initial frequency and f-I gain of initial frequency. On the other hand, cells with higher values of absolute sag, ADP and AHP and resting potential exhibit lower values of PrC2. Lastly, PrC3 may reflect firing of an action potential as the PrC3 is dominated by the values of AP threshold and input resistance. Visualizing the first three PrC of type A and type B L5 cells from control and noise exposed mice showed how the different cell types move further away from each other following a previous noise exposure, validating that the noise exposure has a long-term effect on L5 PC membrane properties.

To separate noise-exposure effects from maturation of membrane properties, such as lowering of input resistance and hyperpolarizing the AP threshold (Kroon et al., 2019) we opted to record from cells of 5-8 weeks old (P38-52) similar to (Gee et al., 2012; Lee et al., 2014) as our initial experiments showed that the resting membrane potential decreased with age for type B PCs, but not type A PCs after the third week of age. Our data supports the work of Joshi and others in which mice aged P24-32 that showed cortico-collicular neurons from A1 to have an average resting membrane potential of −66 mV and corticallosal PCs a resting of −71 mV (Joshi et al., 2015), (our results for mature cells: type A −66 mV and type B −72 mV). Therefore, for the auditory cortex, membrane potential of type A and B cells is indistinguishable in animals younger than P24 (young type B = −64 mV). Interestingly, the lack of distinct sag, ADP and AHP for type B PCs does not appear related to the age/resting potential. Hence, noise trauma experiments in C57 mice should fall within an age window in which cells are mature but before the early onset of age related hearing loss (Winne et al, 2020).

It is generally accepted that loud noise exposure can lead to tinnitus by a cochlear injury triggering peripheral deafferentation and adaptive changes in ascending auditory pathways (Elgoyhen et al., 2015). However, both human and animal studies have shown tinnitus without hearing loss (Langers et al., 2012; Longenecker and Galazyuk, 2016), but few studies separate the two conditions. We have used a tinnitus triggering noise that does not significantly change hearing thresholds (Winne et al., 2020) to show that PCs in the auditory cortex alter their input/output function. Our results further demonstrate that in the absence of underlying cochlear pathology, plastic alterations in higher auditory areas could sustain tinnitus.

In summary, we report differences in membrane properties between type A and type B L5 PCs from >5 weeks old mice and show how parameters spread in the opposite direction 1 week following noise exposure depending on subtype of PCs.

## Appendices

Supplementary table 1, Supplementary table 2, Supplementary table 3

## Acknowledgments

We thank Dr. Bryan Souza for help with matlab scripts for analysis.

