## Supplemental Figure 1, table 1 and table 2 for "Loud noise exposure differentially affects subpopulations of auditory cortex pyramidal cells" (from another version of this preprint: version 1, 2020-08-25)

**SUPPLEMENTARY MATERIAL**

### Supplementary Figure

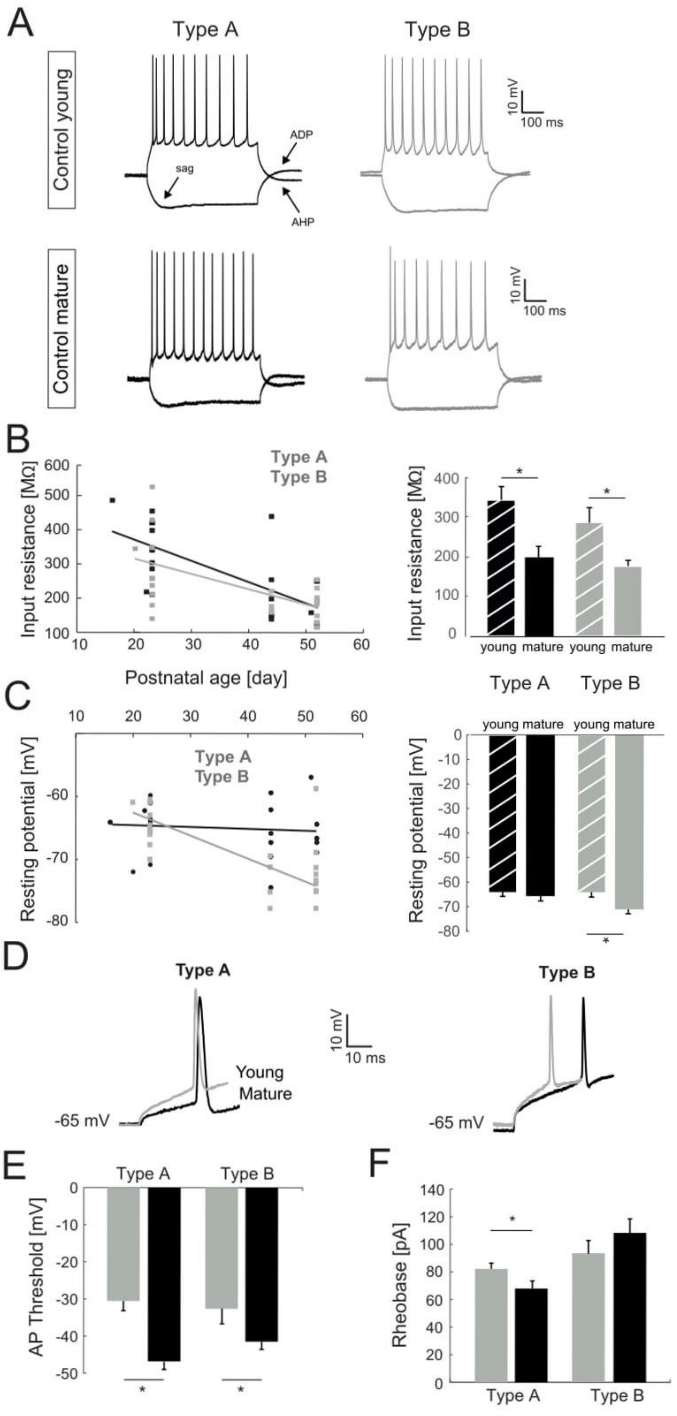


**Supplementary figure 1. Passive and active membrane properties of L5 type A and type B PCs are age dependent.** (A) Representative traces of type A (*black*) and type B PCs (*grey*) from young (P16-23, *up*) and mature (P44-52, *bottom*) mice in response to -100pA and 150pA current injections. (B) Graph showing input resistance versus postnatal age for type A and type B cells (left). Bar graph showing type A and type B PCs decrease input resistance in a more mature age (right). (C) Graph showing resting membrane potential versus postnatal age and group data where type B cells show more hyperpolarized resting membrane potential at older age. (D) Representative traces of the first action potential in response to 150 pA current injection for young and mature type A (*left*) and type B (*right*) L5 PCs of the primary auditory cortex (*right*). (E) Bar graph showing a significant decrease in action potential threshold for both L5 type A ($p=1,6 \times 10^{-5}$) and type B ($p = 0.030$) PCs with age. (F) Bar graph of rheobase (minimum current for generating first action potential) showing significant decrease for type A ($p = 0.014$) PC with age. Error bars - s.e.m., Student's *t*-test, two tailed, equal variances.

### Supplementary Tables

**Supplementary Table 1.** Summary of membrane and AP properties of type A and type B pyramidal cells from juvenile (P16-23) and adult (P44-52) control mice.

| | Type A | | | Type B | | |
|---|---|---|---|---|---|---|
| | Juvenile | Adult | *p* value | Juvenile | Adult | *p* value |
| P (days) | 22.1 ± 0.6 | 47.5 ± 1.2 | *$1 \times 10^{-14}$ **** | 22.7 ± 0.3 | 49.5 ± 1.1 | *$9 \times 10^{-16}$**** |
| $V_{rest}$ (mV) | -64.4 ± 1.1 | -65.7 ± 1.5 | 0.485 | -64.4 ± 1.0 | -72.3 ± 1.4 | *0.0002**** |
| $R_{inp}$ (MΩ) | 361.5 ± 26.2 | 201.9 ± 25.4 | *0.0003**** | 294.5 ± 37.9 | 182.5 ± 11.1 | 0.0004*** |
| Rheobase (pA) | 82.2 ± 3.3 | 68.0 ± 4.3 | *0.014 ** | 93.4 ± 9.5 | 108.4 ± 9.5 | 0.286 |
| Δ Sag (mV) | 2.8 ± 0.4 | 3.4 ± 0.3 | 0.219 | 0.6 ± 0.5 | 0.5 ± 0.1 | 0.972 |
| Δ ADP (mV) | 3.7 ± 0.6 | 3.2 ± 0.9 | 0.645 | 0.5 ± 0.6 | 1.0 ± 0.1 | 0.312 |
| Δ AHP (mV) | 4.7 ± 0.8 | 6.3 ± 2.0 | 0.453 | 2.8 ± 1.0 | 2.0 ± 0.3 | 0.454 |
| $AP_{thres}$ (mV) | -30.5 ± 2.0 | -46.8 ± 1.8 | *$1.6 \times 10^{-5}$ **** | -32.6 ± 3.6 | -41.5 ± 1.9 | *0.030 ** |
| $f_{ini}$ at 150 pA (Hz) | 59.1 ± 6.9 | 64.9 ± 10.9 | 0.651 | 46.0 ±4.2 | 33.2 ± 3.8 | *0.035** |
| $f_{ss}$ at 150 pA (Hz) | 15.7 ± 2.1 | 20.3 ± 1.8 | 0.106 | 17.7 ± 2.3 | 13.3 ± 1.3 | 0.093 |
| Gain_ini (Hz/pA) | 0.28 ± 0.02 | 0.46 ± 0.08 | 0.035 * | 0.26 ±0.04 | 0.38 ± 0.038 | 0.049 * |
| Gain_ss (Hz/pA) | 0.05 ± 0.01 | 0.088 ± 0.04 | 0.357 | 0.05 ± 0.01 | 0.071 ± 0.01 | 0.074 |

P - postnatal; Vrest- resting membrane potential; R_inp - input resistance; Rheobase - minimal current to cause an action potential; Sag - hyperpolarization sag, ADP -afterdepolarization; AHP - afterhyperpolarization; AP_thres - Action potential threshold; f - firing frequency; ini - initial; ss - steady state; f-I gain - frequency over current gain; Student's t-Test, two-tailed, equal variance. Data show Standard Error of the Mean (s.e.m). * $p \leq 0.05$ *

**Supplementary Table 2.** Coefficients of the three first principal components of the data variables

| Variables | PrC1 | PrC2 | PrC3 |
|---|---|---|---|
| Absolute.sag | -0.246 | -0.436 | 0.083 |
| Absolute.ADP | -0.188 | -0.346 | 0.254 |
| Absolute.AHP | -0.213 | -0.264 | -0.316 |
| Resting potential | -0.241 | -0.206 | -0.126 |
| Input resistance | -0.142 | 0.175 | -0.589 |
| AP threshold | 0.204 | 0.036 | -0.525 |
| AP time | 0.263 | 0.186 | 0.015 |
| Rheobase | 0.391 | 0.213 | 0.078 |
| ISI initial | 0.389 | -0.154 | -0.001 |
| Frequency ini. | -0.348 | 0.216 | -0.048 |
| ISI Steady state | 0.369 | -0.271 | -0.087 |
| Frequency SS | -0.305 | 0.385 | 0.14 |
| $F_{ini}$-I gain | -0.115 | 0.204 | -0.27 |
| $F_{ss}$-I gain | -0.046 | 0.287 | 0.287 |
| Relative variability | 0.313 | 0.189 | 0.119 |

ADP- afterdepolarization, AHP- afterhyperpolarization, AP - action potential, AP time - delay to first spike, Rheobase - minimum current to generate an AP using a ramp protocol, ISI - inter spike interval, Ini - initial, SS - steady state, F-I - frequency over current gain

**Supplementary Table 3.** Effects of noise exposure depends on PC type according to multivariate analysis of variance (Manova)

| | MANOVA | Post-hoc (2 way ANOVA) | |
|---|---|---|---|
| | | Contributing variables | p-values |
| **Condition** | Wilk`s trace = 0.77159<br>Num DF =12<br>Den DF = 41<br>p-value = 0.45641 | | |
| **Cell type** | | Absolute sag** | 1.29e-13 |
| | Wilk`s trace = 0.18218 | Absolute AHP** | 4.23e-14 |
| | Num DF =12 | Resting potential | 0.000328 |
| | Den DF = 41 | AP threshold | 0.02153 |
| | p-value = **1.58e-11** | Rheobase*** | 0.004758 |
| | | Fss-I gain* | 0.001531 |
| **Interaction** | Wilk`s trace = 0.56559 | | |
| | Num DF =12 | initial freq.* | 0.02963 |
| | Den DF = 41 | steady-state freq. | 0.03384 |
| | p-value = **0.01071** | | |

Num DF: numerator degrees of freedom; Den DF: denominator degrees of Freedom. Transformed variables before MANOVA: * Ln function; ** Box-Cox transformation; *** inverse function.
